# Quartformer: An Accurate Deep Learning Framework for Phylogenetic Tree Construction

**DOI:** 10.1101/2025.04.05.646867

**Authors:** Junyu Su, Xiyun Jiao

## Abstract

The construction of phylogenetic trees is a fundamental task in evolutionary biology, with the inference of tree topologies being particularly challenging due to the super–exponential growth in the number of possible topologies as the number of species increases. Recent advances in deep learning have offered promising solutions to this challenge, especially approaches that first infer the topologies of all quartets using deep neural networks, and then assemble these quartets into a complete phylogenetic tree using quartet combination algorithms. Building upon these methods, we propose Quartformer, a novel framework that incorporates a sparse attention mechanism to enhance information interaction among quartets derived from the same tree. This allows each quartet to be inferred with richer contextual information, leading to more accurate topology predictions. Experimental results demonstrate that Quartformer outperforms both the original framework and the traditional maximum likelihood method RAxML on simulated datasets. We further validate its improvements on real–biological datasets and theoretically analyze the underlying reasons for its performance gains, which also inform future research directions. Additionally, we systematically evaluate optimization strategies for inference speed and their impact on overall performance. Overall, this work advances the application of deep learning in phylogenetic tree Construction and provides theoretical and practical insights for future methodological developments in the field.

## 1. Background and Related Work

Phylogeny is the science of studying evolutionary relationships among organisms, often focusing on species, individuals, or genes. Its goal is to understand these relationships through the process of evolution. Phylogenetics primarily utilizes molecular data (such as DNA and protein sequences) and morphological data to construct and analyze phylogenetic trees. These trees represent the evolutionary relationships among organisms in a graphical format.Phylogenetics has significant applications across various fields, extending beyond academic research. In biodiversity conservation, understanding evolutionary relationships among species helps identify key species and prioritize their protection. For example, phylogenetic analyses can identify species with critical ecological roles, aiding ecosystem management and balance [15]. In public health and medicine, phylogenetics is widely used to trace the transmission and mutation pathways of pathogens. For instance, during the COVID- 19 pandemic, scientists analyzed the genetic sequences of the virus to trace its origins and transmission routes, enabling the formulation of more effective control strategies [5].

In phylogenetics, the inference of phylogenetic trees is one of the most critical tasks. Reconstructing a phylogenetic tree typically involves two aspects: inferring the tree topology, which determines the evolutionary relationships among species, and estimating branch lengths, which quantify evolutionary distances or times. Tree topology inference is particularly challenging because the number of possible topologies grows super–exponentially with the number of species, making it nearly impossible to exhaustively search for the optimal tree under any given criterion.

Traditional phylogenetic construction methods can be broadly classified into distance matrix–based methods and character–based methods [24]. Distance matrix–based methods calculate pairwise nucleotide sequence substitution distances using Markov chain models to generate a distance matrix. Common models include JC69, K80, HKY85, and GTR, which estimate nucleotide frequencies and substitution rates. To account for varying mutation rates and selection pressures, gamma distributions are often used to adjust rate variation. Once the distance matrix is generated, sequence alignment is no longer required. The main methods for converting the distance matrix into a phylogenetic tree include the least squares (LS) method [4], minimum evolution (ME) method [17], and neighbor joining (NJ) method [18].

Character–based methods are crucial in phylogenetic Construction and include maximum parsimony (MP) [3], maximum likelihood (ML) [7], and Bayesian inference (BI) [16]. These methods directly evaluate character changes at each site in a multiple sequence alignment (MSA) and determine the most suitable tree topology based on specific criteria, such as minimizing changes or maximizing likelihood.

In recent years, machine learning and deep learning techniques have been introduced into phylogenetic tree inference, leading to new methodological advances [11]. Currently, there are three main approaches:

### 1. Accelerating heuristic searches using machine learning models

For example, Azouri et al. (2021) [1] proposed a machine learning method based on random forest regression to predict the tree neighbors most likely to increase the likelihood during phylogenetic tree search, avoiding the costly likelihood calculation for each node and thus significantly accelerating the heuristic tree search process. Building on this, Azouri et al. (2024) [2] further introduced a reinforcement learning–based search strategy aimed at optimizing the overall tree search strategy. Unlike traditional methods that rely on local optimization, this approach uses an agent to learn a global optimization strategy, enabling the search process to converge more efficiently to the optimal tree topology in a shorter amount of time, and demonstrating faster inference speed than traditional methods in longer sequences.

### 2. Predicting distance matrices with deep learning models

This method uses deep learning to infer evolutionary distances, which are then used to reconstruct tree topologies through distance–based methods (such as NJ). For example, Phyloformer (Nesterenko et al., 2022; Nesterenko et al., 2024) is a neural network based on the Transformer architecture that efficiently predicts pairwise evolutionary distances between sequences [12, 13]. These predicted distances are then used in NJ to reconstruct phylogenetic trees. Phyloformer integrates information from the entire MSA through a self attention mechanism, which not only improves the accuracy of distance predictions but also significantly speeds up the inference process. Under the standard LG+GC model, the accuracy of Phyloformer is comparable to that of ML methods, while its speed is improved by two orders of magnitude. Furthermore, under more complex models (such as those considering site dependencies or heterogeneous selective pressures), Phyloformer outperforms other methods.

### 3. Inferring and combining quartet topologies

Deep learning models predict the topologies of all possible quartets (sets of four species), which are then combined to construct a complete tree. Suvorov et al. (2020) [20] and Zou et al. (2020) pioneered this approach [25]. Wang et al. (2023) further developed the Fusang framework, optimizing the data generation and model inference processes, and demonstrated the effectiveness of the method through extensive experiments [22]. In addition, Tang et al. (2024) [21] proposed a novel symmetry– preserving neural network architecture that significantly improves model robustness under long–branch attraction conditions. Yakici (2023) [23] introduced a framework based on multi–layer perceptrons (MLPs), using a Bag–of–Words–like embedding of site patterns to scale phylogenetic inference to genome–level datasets.

This study is based on the third approach mentioned above, and introduces an innovative framework by incorporating a sparse attention module between the deep learning model for quartet classification and the quartet combination algorithm. This module enables the hidden vectors of different quartets in the same phylogenetic tree to interact, thereby improving the accuracy of quartet topology prediction and, in turn, enhancing the overall accuracy of phylogenetic tree inference.

The structure of this paper is organized as follows: Section 2 describes the hardware configuration of the experimental platform and the methods used for generating simulated data. Section 3 provides a detailed explanation of the structure and design principles of the Quartformer framework. Section 4 first presents the framework’s performance on simulated datasets, including accuracy, inference speed, and optimization strategies, and then evaluates the model’s practical applicability on real–biological datasets. Section 5 offers a theoretical analysis of Quartformer’s advantages over the original framework in the quartet prediction task. Finally, Section 6 summarizes the main contributions of this study and discusses potential directions for future improvements of deep learning methods in phylogenetic tree construction.

## 2. Experimental Platform and Simulated Data Generation

All experiments were conducted on a workstation equipped with an Intel Core i9-14900K processor (24 cores), 128 GB DDR5 4200 MHz memory, and an NVIDIA GeForce RTX 4090 GPU (24 GB). The data simulation process consists of two main steps: (1) generating phylogenetic trees and (2) using the generated trees to simulate MSAs for all species using the Indelible software [8].

To evaluate the model on simulated datasets, we adopted the simulation method described in the Fusang framework [22, 28]. The topologies of phylogenetic trees were randomly generated using the ETE3 Python library [9], which allows for the generation of random tree structures with a given number of leaves (see Figure 2A). Branch lengths were sampled from either a gamma distribution or a uniform distribution (Figure 2B) to define relative branch lengths. To ensure biological relevance, Fusang analyzed the pairwise divergence distributions between leaf nodes from real datasets and used this distribution to rescale internal branch lengths, resulting in phylogenetic trees that more closely resemble realistic evolutionary processes. Simulated datasets were categorized into six variants based on the branch length distribution and sequence types: SG, SU, NG, NU, CG, and CU. Here, C, N, and S represent coding sequences, non-coding sequences, and their mixtures, respectively, while U and G indicate the use of uniform and gamma distributions for sampling branch lengths.

**Figure 1:**
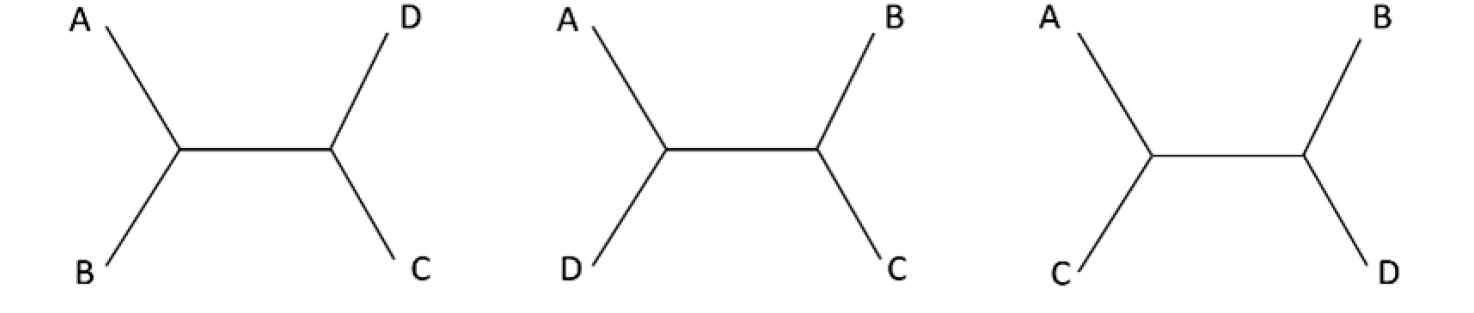
The three possible topologies of a quartet.

**Figure 2:**
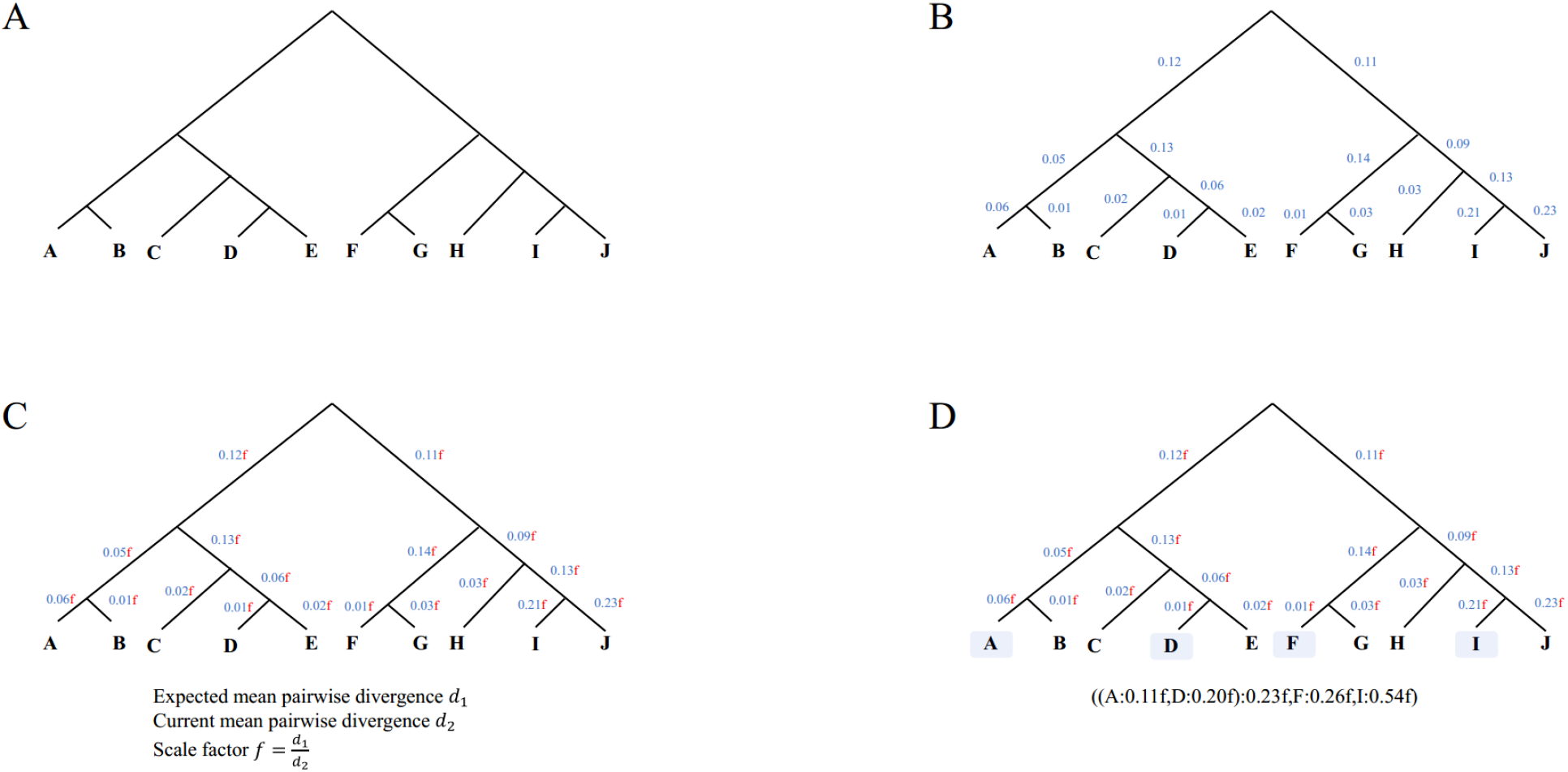
Fusang framework simulation process. [22]

Previous studies have shown that deep learning models are relatively insensitive to the choice of evolutionary model. To verify this property, we generated training data using the GTR evolutionary model and trained a CNN model based on the architecture proposed by Suvorov [20]. During testing, MSAs were simulated under five different evolutionary models: JC69, TIM, TIMef (equal base frequencies), GTR, and UNREST. The experimental results (see Figure 3) confirmed the robustness of the model to variations in evolutionary assumptions. To ensure the simplicity of the simulated dataset and enable fair comparison— especially with methods such as ML, which are more sensitive to model assumptions—we fixed the simulation model to GTR for all datasets. Furthermore, while deep learning models can extract evolutionary information from insertion–deletion (Indel) patterns in MSAs, ML methods typically treat such symbols as ambiguous. As a result, deep learning models often significantly outperform ML-based methods in terms of topology inference accuracy when the MSA includes Indels. To ensure fairness in comparison, no Indels were included in our simulated datasets.

**Figure 3:**
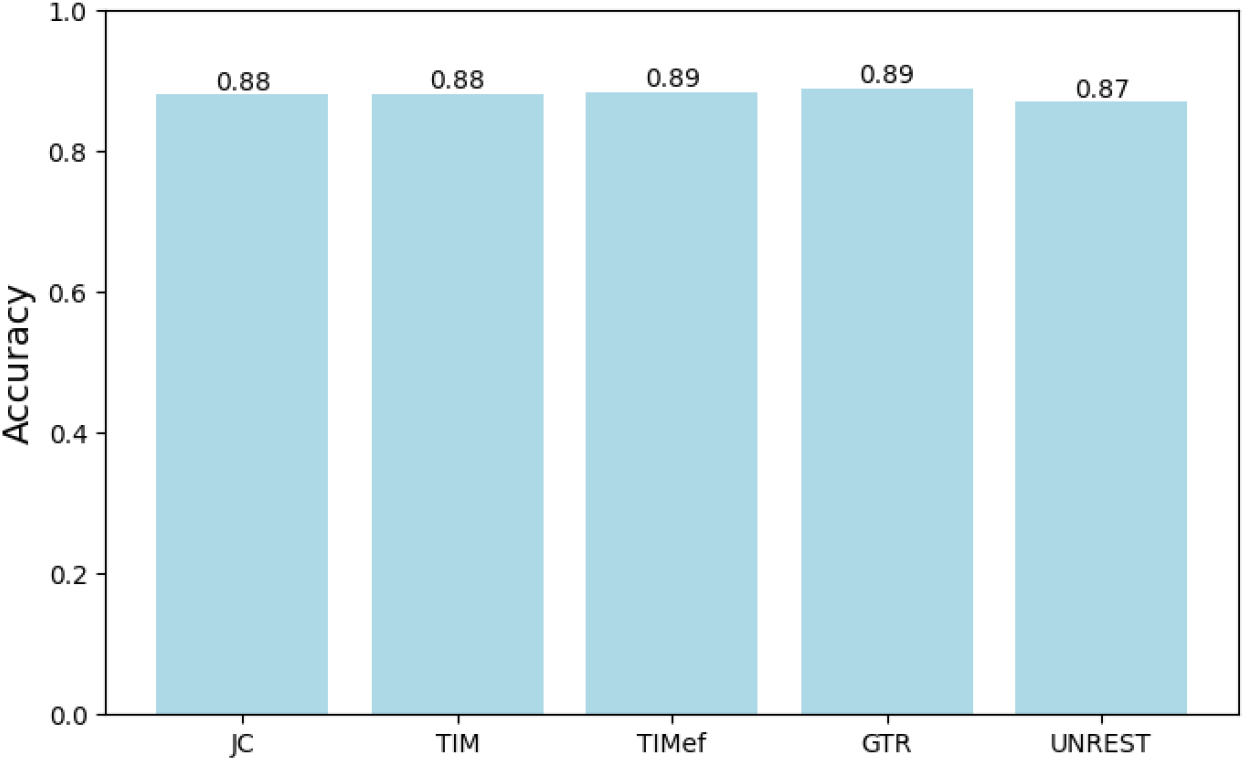
Performance of CNN under different evolutionary models.

**Figure 4:**
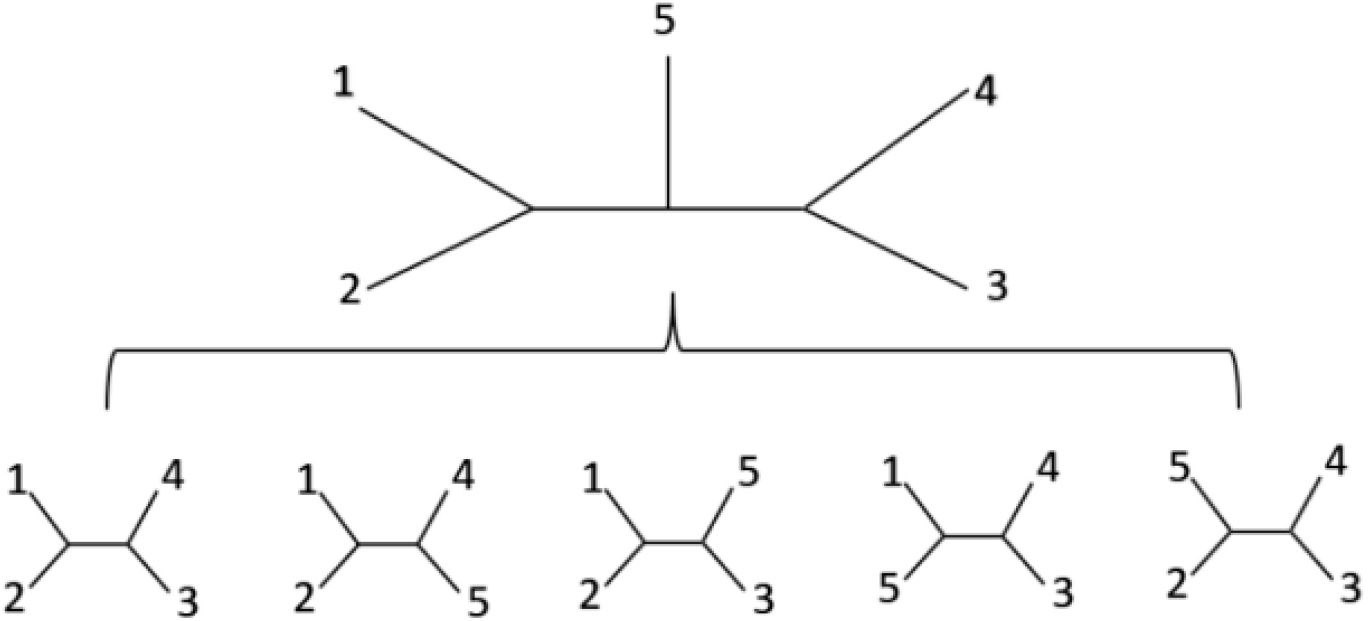
Illustration of inducing a quartet topology from a phylogenetic tree with five species (1, 2, 3, 4, 5).

The simulation parameters for Indelible followed those used in Suvorov’s study. Specifically, to model rate heterogeneity along the sequence, we sampled the shape parameter of the continuous gamma distribution from the interval *U* (0, 4). The proportion of invariable sites (pinv) in each MSA was sampled from *U* (0, 1). Note that the expected number of substitutions per site (i.e., branch length) was specified only for variable sites. For instance, when pinv = 0.5, the specified branch lengths apply only to the 50% of sites that are variable, while the remaining invariable sites have zero branch length. Substitution rate parameters were sampled from *U* (0, 3), and nucleotide frequencies were drawn from a four–category Dirichlet distribution with concentration parameter *α* = 30.

For phylogenetic trees with more than four species, quartet MSAs were induced and their corresponding topologies determined. Each MSA was converted into a numeric matrix using the encoding {A: 0, C: 1, T: 2, G: 3}, resulting in a 4 *×* 1000 input matrix for each quartet. The four DNA sequences were assigned labels 1 through 4. The quartet topology was then one–hot encoded as follows: ((1, 2), 3, 4) : [1, 0, 0], ((1, 3), 2, 4) : [0, 1, 0], ((1, 4), 2, 3) : [0, 0, 1]. These encoded matrices and their corresponding labels formed the input–target pairs used to train the deep learning model. Since all deep learning components in the Quartformer framework are designed to perform classification over quartet topologies, we adopted the cross–entropy loss function as the training objective. For the three–class classification task, the cross–entropy loss is defined as:

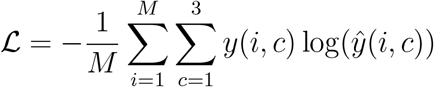

where *M* is the number of samples, *c* is the class index, *y*(*i, c*) is the one–hot true label for the *i*th sample in class *c*, and *ŷ*(*i, c*) is the predicted probability for that class.

## 3. Baseline and QuartFromer Framework

First, we introduce the baseline framework, which consists of two main components. The first component is the deep learning model, which focuses on predicting the quartet topology. Specifically, the model takes the DNA sequences of four species as input and outputs a probability distribution of which of the three possible quartet topologies these species belong to. The second component is the quartet combination algorithm. The second component takes the quartet topology predicted by the deep learning model, which is a probability-weighted topology based on the model’s output distribution, as input and generates a complete phylogenetic tree that includes all species. These two components are independent and solve different sub–tasks.

In this study, we selected the CNN model proposed by Suvorov and the wQFM algorithm [10] as the baseline framework. The CNN model consists of a feature extraction module and a classification head. The feature extraction module comprises eight convolutional and pooling layers followed by a linear mapping layer, which encodes the MSA of a quartet (with a length of 1000) into a 128-dimensional feature vector. The classification head then maps this feature vector into a three–class probability distribution via a fully connected layer. The wQFM algorithm serves as the quartet combination module, which integrates the predicted topologies of all quartets to construct the final phylogenetic tree. These two methods represent excellent techniques in their respective domains and have demonstrated higher accuracy than RAxML [19] in simulation experiments. Based on these two core components, we designed the overall research workflow, as illustrated in Figure 5. This workflow achieves modular decoupling of tasks and fully leverages the expressive power of the deep learning model in classification as well as the optimization capability of the combination algorithm. The figure demonstrates how quartet classification and combination algorithms work together to accomplish phylogenetic tree construction.

**Figure 5:**
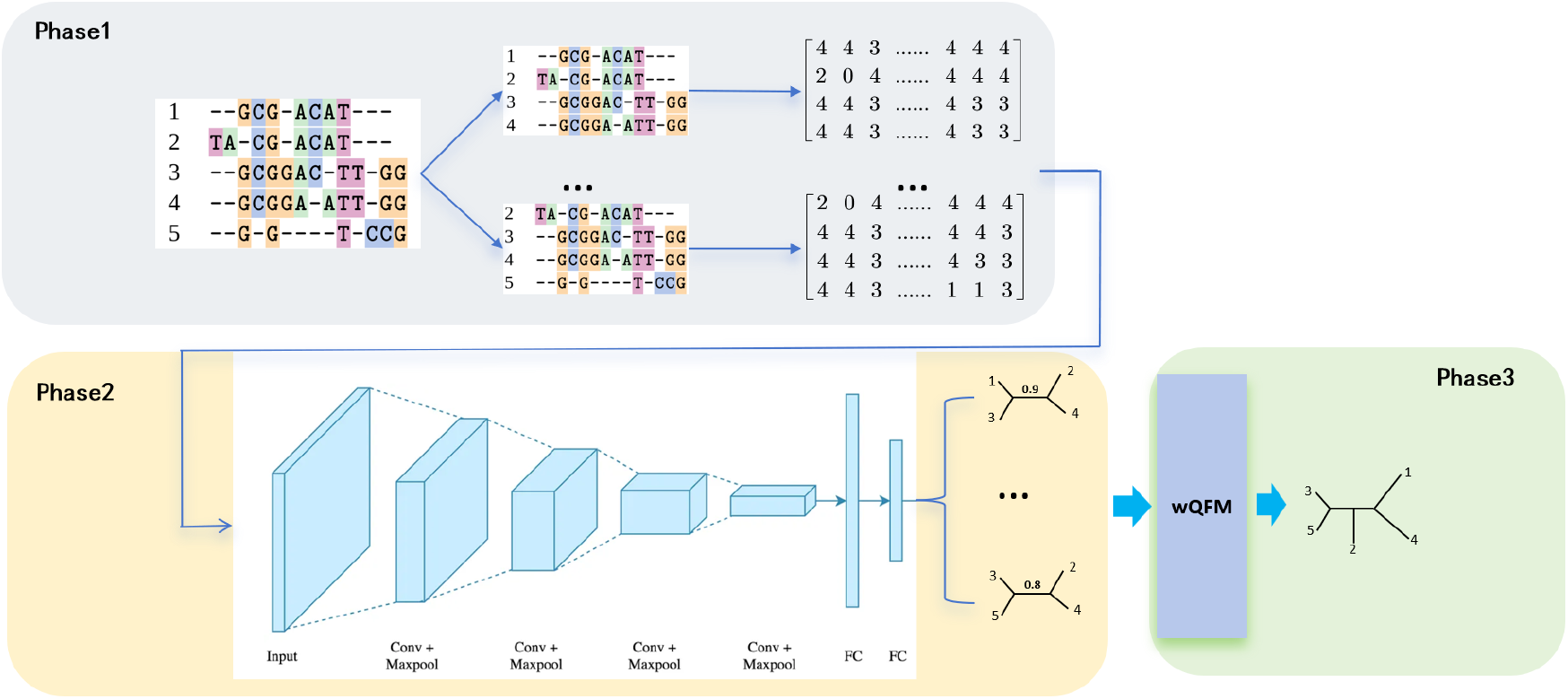
Workflow of the baseline framework.

From this framework, it is clear that factors affecting the accuracy of the final phylogenetic tree topology include the classification accuracy of the CNN and the search ability and robustness of the combination algorithm. To quantify the impact of quartet classification accuracy on the final tree Construction, we generated 100 phylogenetic trees for 10, 20, 30, 40, and 50 species. For each tree, we induced all possible quartets and artificially modified their topologies to simulate CNN classification accuracies of 0.95, 0.85, 0.75, and 0.65. We used the normalized Robinson–Foulds (RF) distance to measure the difference between the wQFM–reconstructed tree and the true tree. The results, shown in Figure 6, met our expectations.

**Figure 6:**
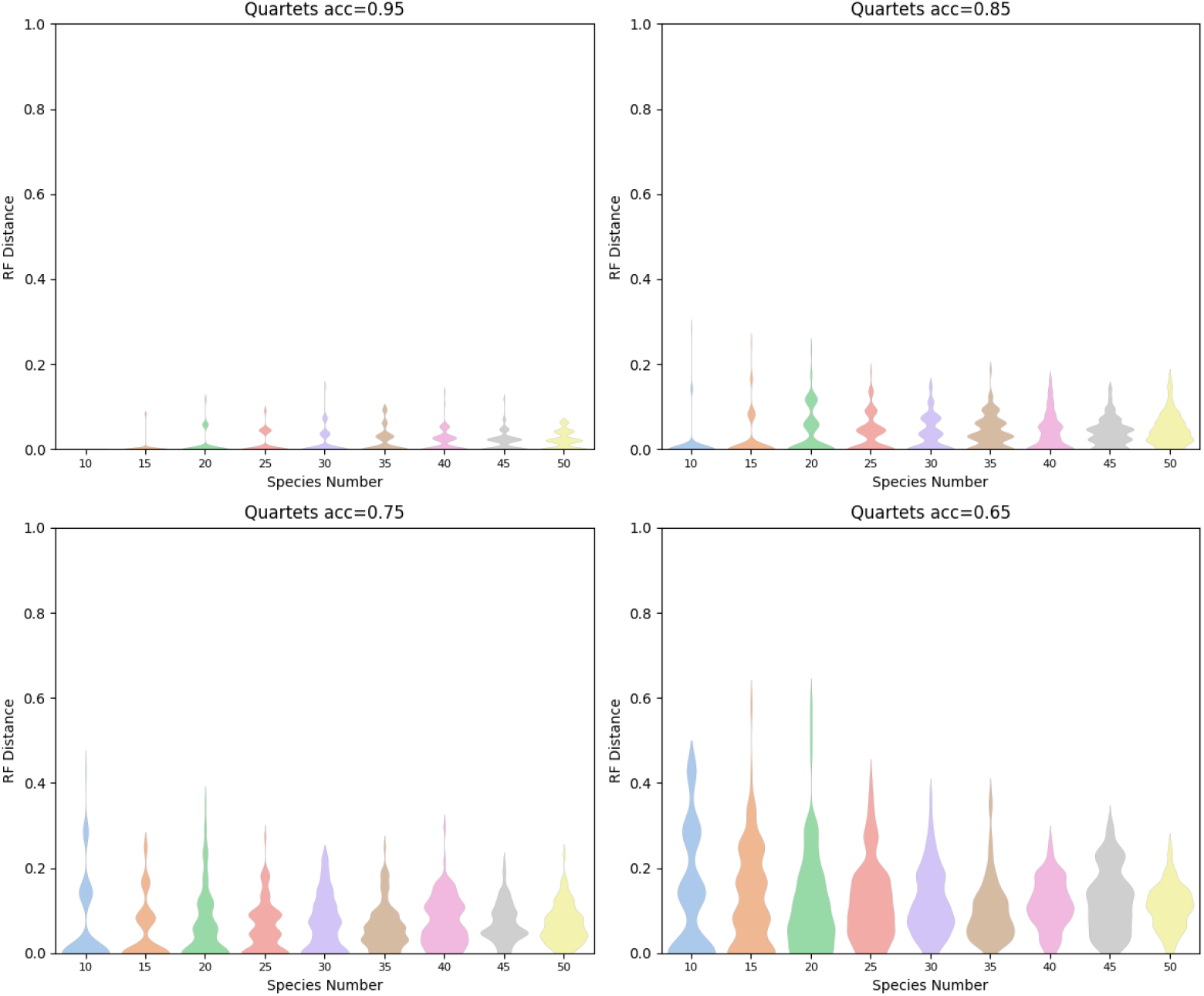
Normalized RF distance between the reconstructed tree and the true tree for different quartet classification accuracies (0.95, 0.85, 0.75, and 0.65) with varying numbers of species (10-50).

**Figure 7:**
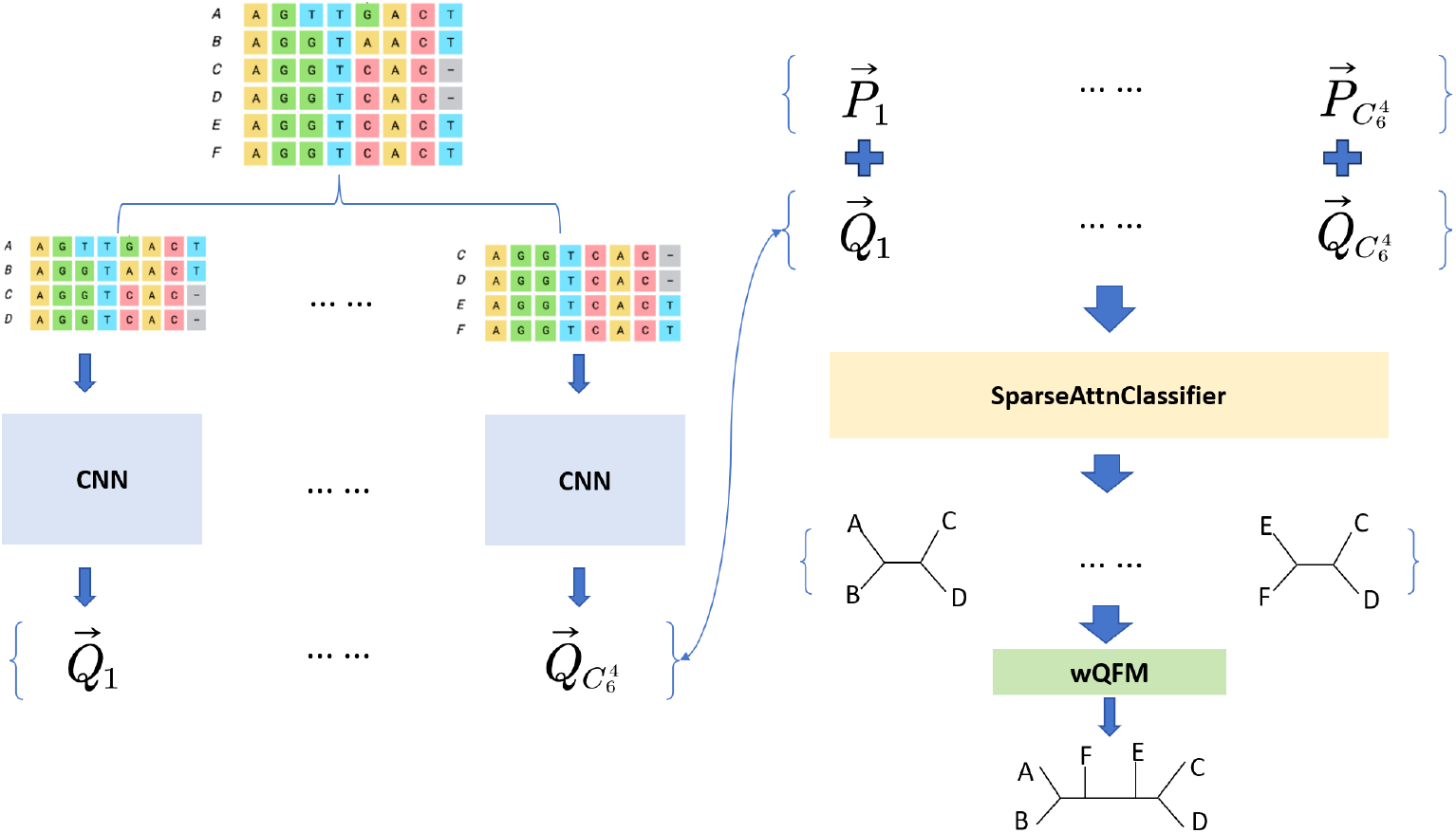
Workflow of the Quartformer framework.

We now introduce the Quartformer framework, which consists of three core components: the feature extraction module, the sparse attention module, and the quartet combination module. The feature extraction module uses the CNN model from the baseline framework with the classification head removed. This module maps the MSA of each quartet into a 128-dimensional vector. The second module first concatenates a 128-dimensional vector to each quartet’s vector, representing the taxa included in the quartet. It then performs sparse attention calculations and fusion updates between the quartet vectors at specified positions, finally mapping the result into a three–class probability distribution. The third component is the quartet combination algorithm, wQFM, which treats the topology probability distributions of all quartets as weights and generates the final output. The Quartformer framework is highly flexible and can be continuously updated with different modules in future work.

Before detailing the calculation process of the sparse attention module, we explain the motivation behind the design. Specifically, the quartets induced from the same phylogenetic tree are not fully independent. For example, a phylogenetic tree with five species can induce five quartets, as shown in Figure 8. If the second quartet, ((1, 2), (4, 5)), is removed, we can still correctly infer the complete phylogenetic tree that includes all species. Moreover, if the second quartet is transformed into another topology, it will contradict the other four correct quartets. This suggests a pattern: the quartet topologies derived from the same phylogenetic tree are interdependent. Based on this observation, we chose the attention module to learn this interdependency. Specifically, if the classification model can predict most of the quartet topologies correctly, the majority of correct quartet topologies can help correct the inconsistent ones.Considering that the correlations between quartets vary, we restrict each attention layer to calculate attention weights only for quartets that share three taxa. This ensures that attention is computed only for relevant subsets of quartets. Additionally, to ensure that information exchange between quartets is both global and hierarchical, we design the attention mechanism with four layers. The final network architecture is a sparse multi-layer attention network based on the shared taxa model.

**Figure 8:**
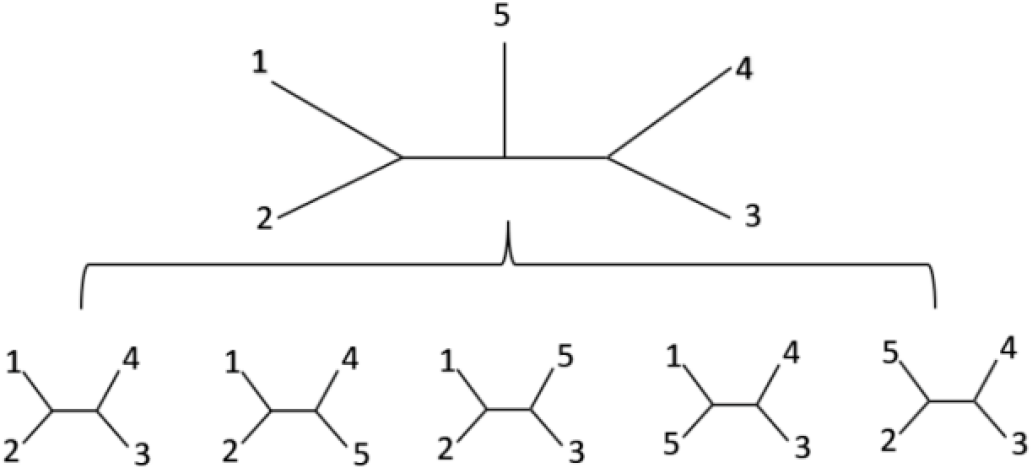
Illustration of inducing a quartet topology from a phylogenetic tree with five species (1, 2, 3, 4, 5).

**Figure 9:**
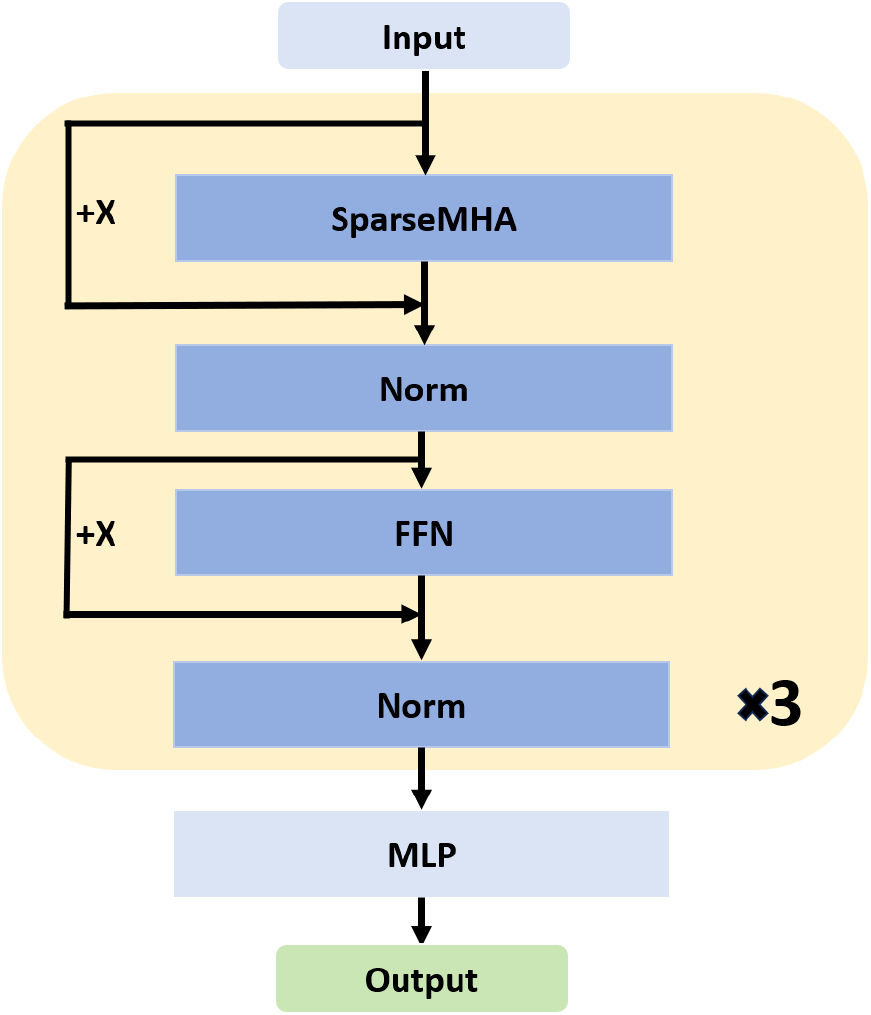
Calculation process of the second module.

The overall computation process of the sparse attention module is described as follows. All quartet embeddings produced by the CNN model have shape (*N*, 128), where *N* is the number of quartets. Each embedding is concatenated with a 128-dimensional identification vector. Each identification vector is divided into *n* segments (where *n* is the total number of taxa). For any given quartet, the segments corresponding to its four taxa are set to 1, and all remaining positions are set to 0. For example, if a quartet contains taxa 1, 2, 3, and 4, the first four segments of the identification vector are set to 1 and the rest to 0. This results in a final 256-dimensional input vector (128 from the encoder and 128 from the identification vector), forming a matrix **X** *∈* ℝ^*N×*256^. To ensure proper alignment between sequence order and identification vector, the four DNA sequences in each quartet MSA must be sorted in ascending order by taxa labels. Each Transformer layer consists of a sparse self attention operation followed by a feed–forward network. First, **X** is linearly projected to obtain the query, key, and value matrices as **Q** = **XW**_*Q*_, **K** = **XW**_*K*_, **V** = **XW**_*V*_, where **W**_*Q*_, **W**_*K*_, **W**_*V*_ ∈ ℝ^256×256^ are learnable parameter matrices. A sparse mask matrix **M** *∈ {*0, *−∞}*^*N×N*^ is then defined based on shared taxa information: if two quartets share at least three taxa, **M**_*ij*_ = 0; otherwise, **M**_*ij*_ = *−∞*. The masked attention weights are computed as

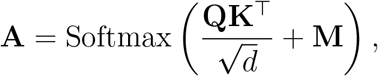

where Softmax is applied row-wise. The attention output is computed as **O** = **AV**, and a residual connection with layer normalization is applied to obtain the intermediate output 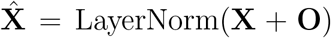. The feed–forward network then projects 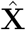 into a higher– dimensional space of size *h*, applies a ReLU activation, and projects it back to the original 256-dimensional space. Here, *h* is a tunable hidden dimension hyperparameter. Let **W**_1_ ∈ ℝ^256*×h*^, **W**_2_ ∈ ℝ^*h×*256^, **b**_1_ ∈ ℝ^*h*^, **b**_2_ ∈ ℝ^256^ be the learnable projection weights and bias vectors. Given that 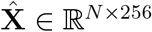, the product 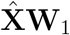 yields a matrix in ℝ^*N ×h*^. The bias **b**_1_ is broadcast across all rows to match the shape (*N × h*) before addition. Similarly, after the second linear transformation, **b**_2_ is broadcast across rows to match (*N ×* 256). The full feed–forward computation is thus given by

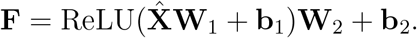

A second residual connection and layer normalization are applied as 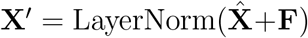. After stacking multiple Transformer layers, the final output embeddings **X**′ ∈ ℝ^*N ×*256^ are passed through a linear classifier to predict the three–class quartet topology distribution as

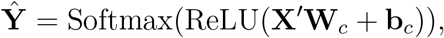

where **W**_*c*_ ∈ ℝ^256*×*3^, **b**_*c*_ ∈ ℝ^3^. Since the result of **X**^*′*^**W**_*c*_ has shape (*N ×* 3), the bias **b**_*c*_ is broadcast along the rows and added element–wise to obtain the final prediction **Ŷ** ∈ ℝ^*N ×*3^, where each row encodes the predicted probability distribution over the three possible unrooted quartet topologies.

In summary, the sparse attention module effectively captures inter–quartet relationships while avoiding redundant computations between unrelated quartets. By stacking multiple layers, this module integrates both local and global information, thereby significantly enhancing the final representational capacity. In terms of implementation, our sparse attention mechanism is inspired by the design of FlashAttention V2 [6]. It is important to note that the sparsity pattern is determined by the number of shared taxa between quartets. As the number of quartets increases, directly storing a sparse mask matrix of size number of quartets *×* number of quartets on GPU becomes infeasible due to memory constraints. To address this challenge, we adopt a block–wise dynamic masking strategy during FlashAttention’s block–wise computation. Specifically, within each attention block, the corresponding mask is dynamically computed on–the–fly during the computation, without the need to store the entire sparse mask matrix in advance. This fused operator design significantly reduces memory consumption. For instance, when the number of taxa reaches 36, compared to the naive method that requires storing the full sparse mask matrix and the standard FlashAttention method that also requires storing the full sparse mask matrix, our sparse attention implementation reduces memory usage to approximately 1*/*140 and 1*/*70, respectively. The detailed forward and backward computation algorithms for the sparse attention module are provided in the GitHub repository.

The training of Quartformer is designed as a two–stage process. In the first stage, we train the CNN feature extraction module to extract informative representations from quartet MSAs. To achieve this, a classification head is temporarily appended to the CNN feature extraction, and the combined model is trained as a standard three–class classifier using labeled quartet MSAs. Given a quartet MSA *x*, the feature extraction produces a feature vector Encoder(*x*) ∈ ℝ^128^, which is then passed through the classification head composed of a linear layer followed by ReLU and Softmax activations. The output probability distribution over the three possible topologies is computed as

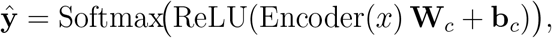

where **W**_*c*_ ∈ ℝ^128*×*3^ and **b**_*c*_ ∈ ℝ^3^ are the learnable weights and biases of the classification head. The output **ŷ** ∈ ℝ^3^ represents the predicted probability distribution over the three unrooted quartet topologies. In the second stage, we use the pretrained CNN feature extraction to generate embeddings for all quartet MSAs in the training set, effectively replacing raw MSA inputs with feature vectors. Then each quartet embedding is further concatenated with an identification vector **p**, resulting in:

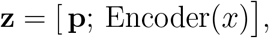

where [*·*;*·*] denotes vector concatenation. The resulting embeddings **z** form the input to the sparse attention module, which models inter–quartet dependencies and outputs topology probabilities for each quartet.

## 4 Results

### 4.1 Experiments with Simulated Data

This section presents a comparison of the accuracy between the Quartformer framework, the baseline framework, and RAxML on simulated datasets. The evaluation is carried out from three perspectives: 1) the accuracy of deep learning models for quartet classification in both frameworks; 2) the accuracy of reconstructing the complete phylogenetic tree topology using the three methods; 3) the effect of increasing DNA sequence length on the quartet classification accuracy of the two deep learning models and RAxML.

#### Quartet Classification Accuracy and Topology Accuracy

As discussed earlier, we quantified the direct impact of deep learning model accuracy for quartet prediction on the output topology accuracy of the wQFM combination algorithm. To comprehensively evaluate the performance of deep learning components in the two frameworks, we used eight test datasets with species numbers ranging from 8 to 36. Each dataset was further divided into six subdatasets based on branch length sampling methods (SG, SU, NG, NU, CG, CU), each containing 100 trees, with a fixed DNA length of 1000. For each tree, we induced the MSA matrix of all quartets, which served as the input to the deep learning model, and the model’s topology prediction for all quartets was the output. The model’s performance was measured by the ratio of “correct classifications/total quartets.” For RAxML’s accuracy, we used the same standard, but due to the slower computation of RAxML, we only sampled 50 quartets for each tree.

Figure 10 (left) shows the average accuracy of the baseline, the Quartformer, and RAxML across all datasets. It is evident that the deep learning methods significantly outperform RAxML in quartet classification, and further improvements are observed when attention layers are added to the deep learning models. The chart also shows the proportion of samples with accuracy improvements after adding the attention layer for each dataset. The results indicate that the model’s predictive ability was enhanced in most cases after adding the attention layer, further validating the robustness of the Quartformer approach. Additionally, the chart on the right shows the distribution of accuracy improvements, with many samples exhibiting accuracy increases of 0.1 or even 0.2, highlighting the significance of the improvements.

**Figure 10:**
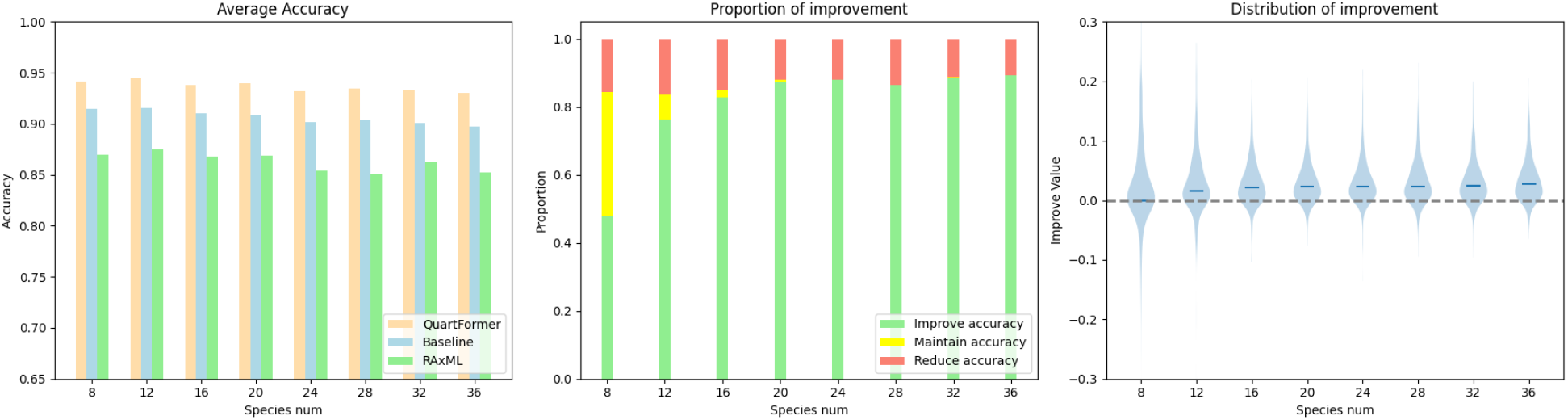
Average accuracy for quartet classification across all datasets.

**Figure 11:**
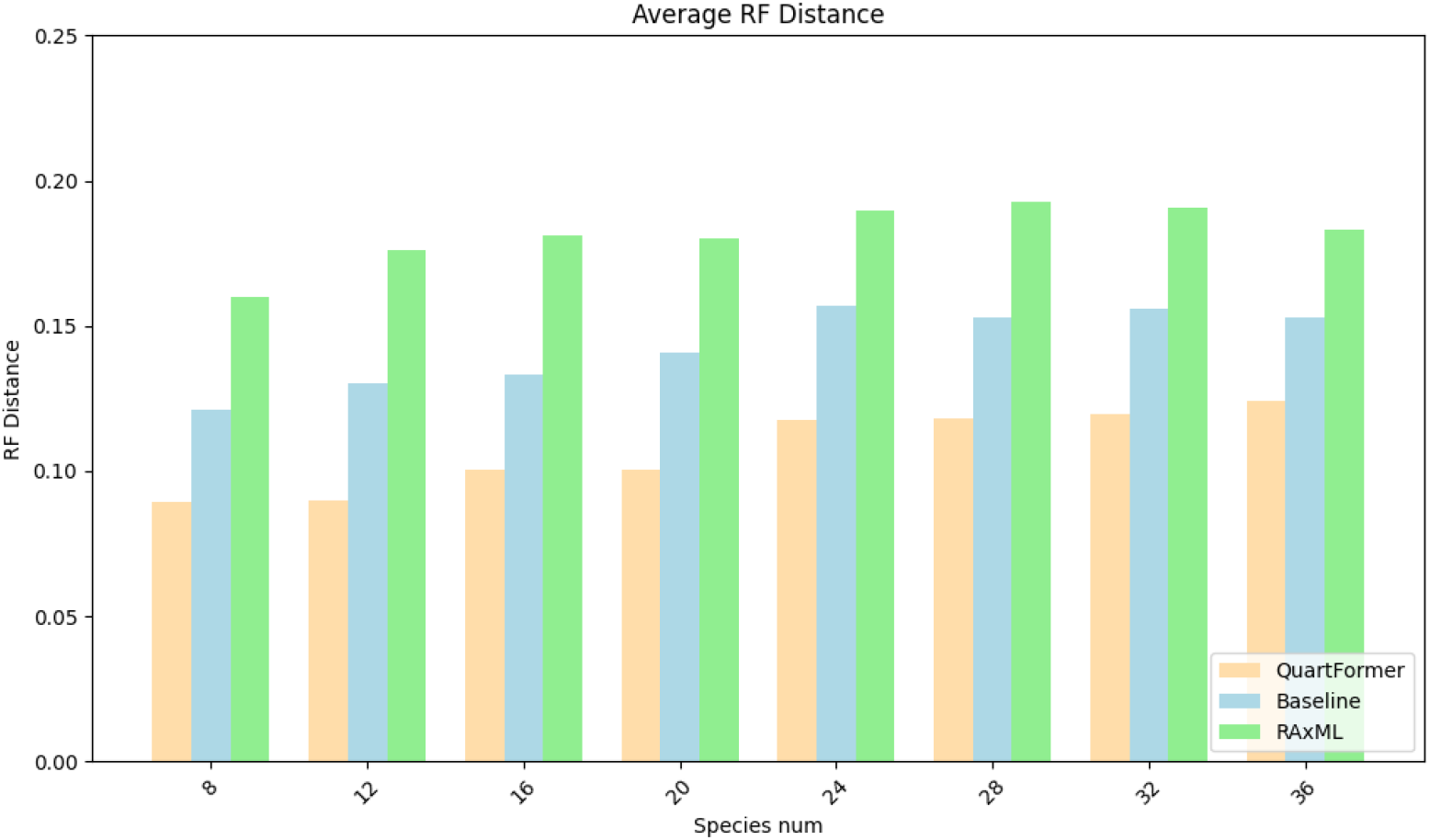
Normalized RF Distance comparison for phylogenetic tree construction across all datasets.

**Figure 12:**
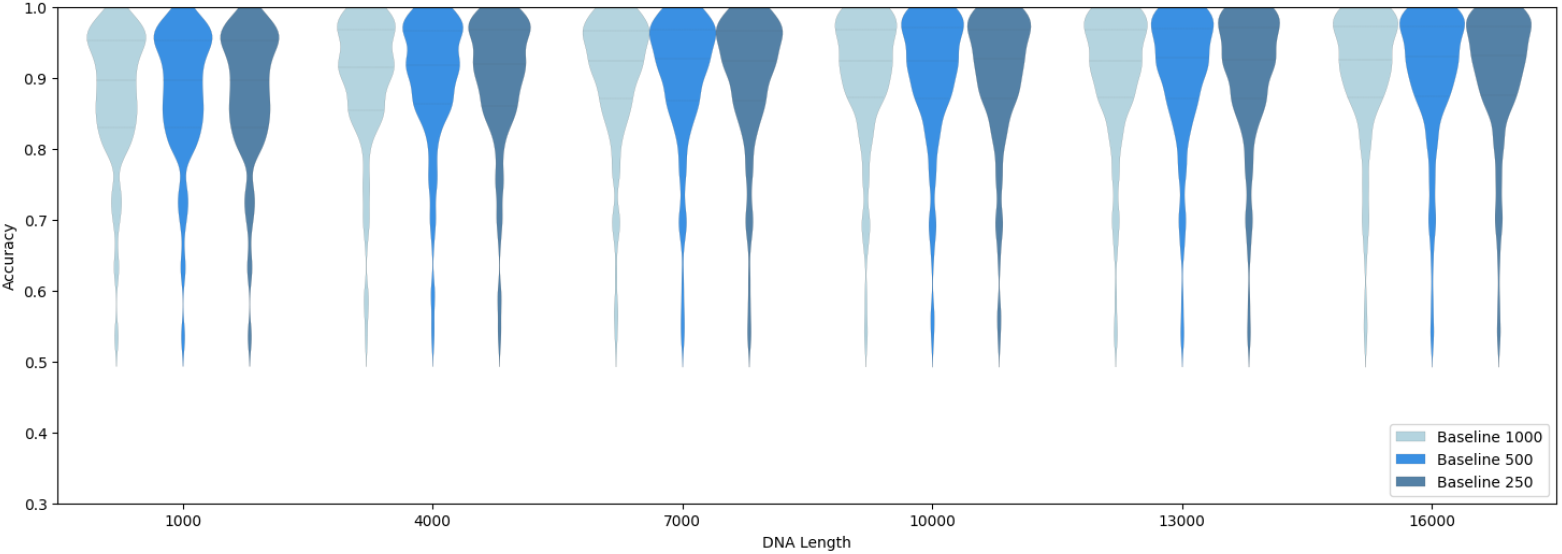
Effect of sliding window strides on baseline accuracy for varying DNA lengths.

**Figure 13:**
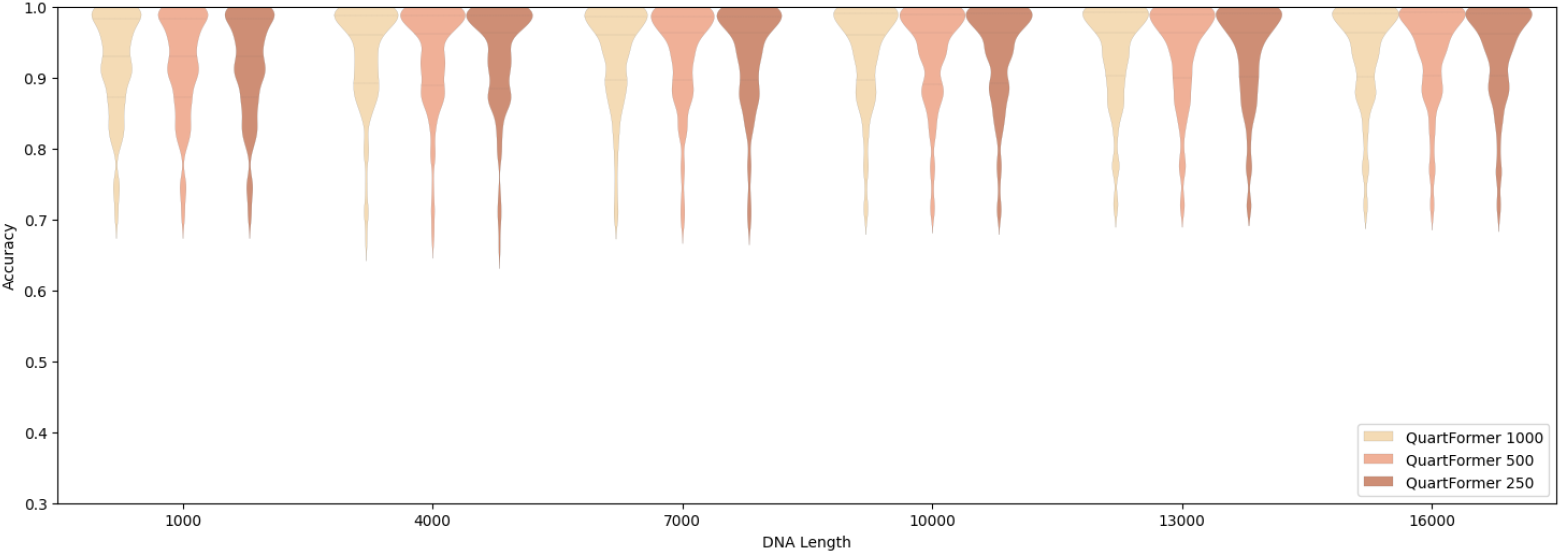
Effect of sliding window strides on Quartformer accuracy for varying DNA lengths.

Subsequently, we collected the predictions of the deep learning models for all quartets of each tree across the same eight datasets and assembled them into complete phylogenetic trees using the wQFM algorithm. We used the normalized RF distance to evaluate the performance of both deep learning frameworks and RAxML on each sample. The figure shows the average normalized RF distances across all samples in the eight datasets. The Quartformer framework achieved the best performance, followed by the baseline framework, while RAxML exhibited comparatively weaker results. These experimental findings demonstrate that the integration of a sparse attention model significantly enhances the accuracy of phylogenetic topology prediction.

#### Comparison of performance with increasing DNA length

In practical applications, DNA sequence lengths often exceed 1000 bases. Therefore, it is important to examine whether the accuracy of deep learning models improves as DNA length increases. In this experiment, we fixed the number of species in the phylogenetic tree to 12 and sampled branch lengths uniformly from six different modes (SG, SU, NG, NU, CG, CU), resulting in 120 trees in total. The simulated MSA length was set to 16000 bases, and then we created six datasets by truncating the MSA to the first 1000, 4000, 7000, and so on base pairs. The accuracy metrics for the samples followed the same evaluation method as described in Section 4.1 for quartet classification. Since there are only 495 quartets in total for the 12 species, RAxML also needed to predict all quartets to calculate its accuracy.

Since the deep learning model requires a fixed MSA input length of 1000, we adopt a sliding window strategy to process longer DNA sequences. Specifically, sequences exceeding 1000 bases are segmented into windows of 1000 bases according to a predefined step size (250, 500, or 1000). The step size determines the overlap between adjacent windows: a step size of 1000 results in non-overlapping windows, while step sizes of 250 or 500 introduce varying degrees of overlap. This approach ensures that the input data meets the model’s length requirements while enabling efficient processing of longer sequences. We evaluated the impact of different step sizes on accuracy and found that window overlap had no significant effect. Therefore, to reduce computational cost, we selected a step size of 1000 for subsequent experiments.

Figure 14 shows the performance comparison of the three methods across datasets of varying lengths. All three methods showed improvements with increasing DNA length. Among the methods, RAxML exhibited the most substantial improvement in accuracy, but it still performed poorly in a few cases. Compared to the baseline framework, the Quartformer framework demonstrated a faster rate of accuracy improvement.

**Figure 14:**
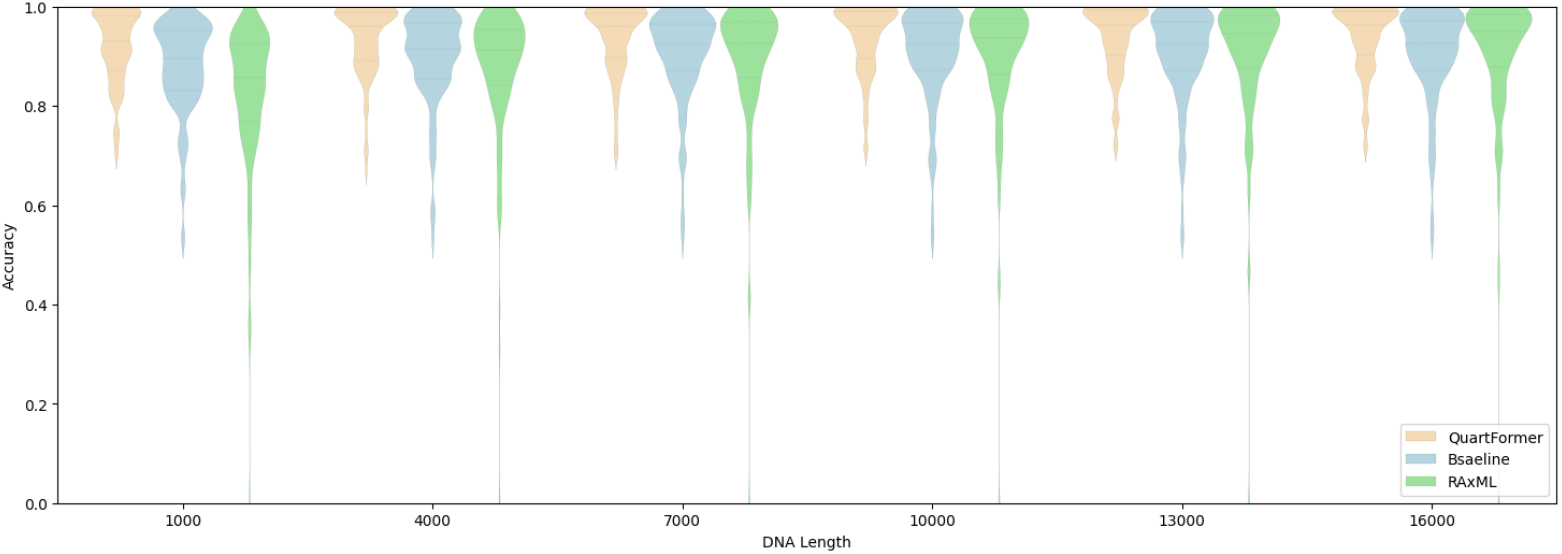
Performance comparison of RAxML baseline and Quartformer across different DNA lengths.

#### Inference Speed Benchmarking and Optimization Strategy

In this subsection, we focus on testing the inference speed of the Quartformer framework, as the speed of inference directly impacts its practical applicability. The computational cost of the Quartformer framework can be divided into three main parts: 1) mapping the MSAs of all quartets into vector representations using the CNN feature extraction module, 2) sparse attention layer computation, and 3) the combination overhead of the wQFM algorithm. It is important to note that the number of quartets grows at a rate of *O*(*n*^4^) with the increase in the total number of species. For example, a phylogenetic tree with 36 species can induce 58,905 quartets, while a tree with 12 species only has 485 quartets. This polynomial growth in the number of quartets means that, as the number of species increases, mapping the MSAs of all quartets into vector representations becomes a bottleneck in terms of computational speed.

To address this issue, we propose an optimization strategy to reduce computational load. Since existing quartet combination algorithms typically do not require the prediction of all possible quartets, we can ignore part of the quartets. Figure 15 presents the inference time of Quartformer under different numbers of species and DNA sequence lengths and compares the time required for predicting all quartets versus ignoring 25%, 50%, and 75% of the quartets (i.e., predicting only 75%, 50%, and 25% of the quartets, respectively, as indicated by 0.75, 0.5, and 0.25 in the figure).

**Figure 15:**
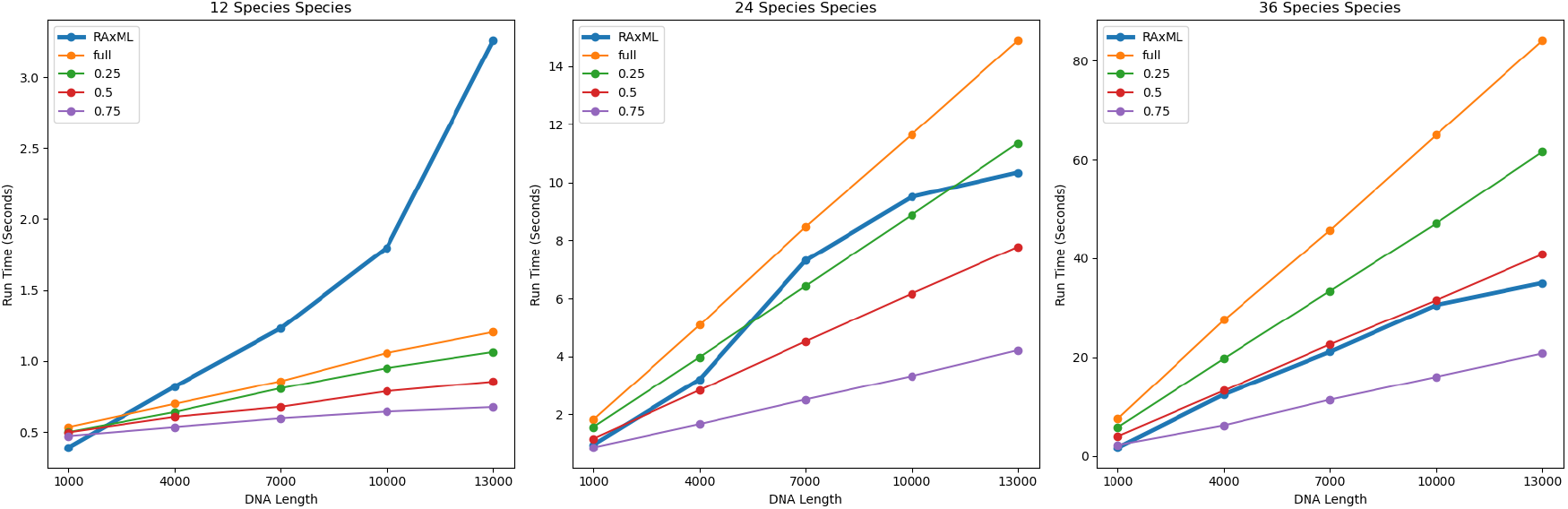
Run time comparison for different species and DNA lengths with varying strategies.

Next, we investigate whether ignoring 25% to 75% of quartets affects the accuracy of both the deep learning components and the overall framework. To this end, we conducted experiments on datasets containing 12, 24, and 36 species, with a fixed DNA sequence length of 1000 bp. Branch length sampling methods were uniformly drawn from the SG to CU simulation strategies. The evaluation criteria and sample definitions follow Section 3.2, where we compared the prediction accuracy of the deep learning model and the final accuracy of the phylogenetic trees reconstructed by wQFM. Figure 16 shows the prediction accuracy of the Quartformer model under different quartet sampling ratios. From the results, it can be seen that although ignoring 75% of quartets leads to a slight drop in accuracy compared to the full prediction strategy, the overall performance still surpasses that of the baseline framework.

**Figure 16:**
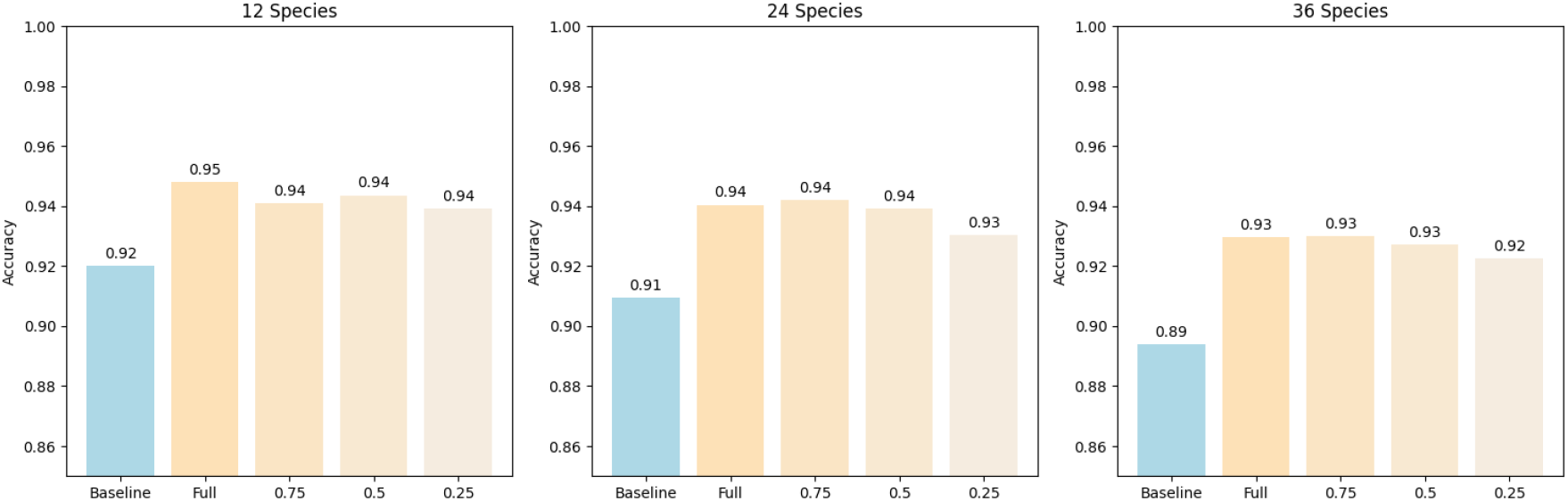
Prediction accuracy of Quartformer under different quartet sampling ratios.

Subsequently, we fed the predicted quartet topologies into the wQFM algorithm to reconstruct the phylogenetic trees, and the results are shown in Figure 17. From the reconstructed trees with 12 to 36 species, we observe that appropriately ignoring a portion of quartets effectively reduces computational cost while maintaining high phylogenetic inference accuracy.

**Figure 17:**
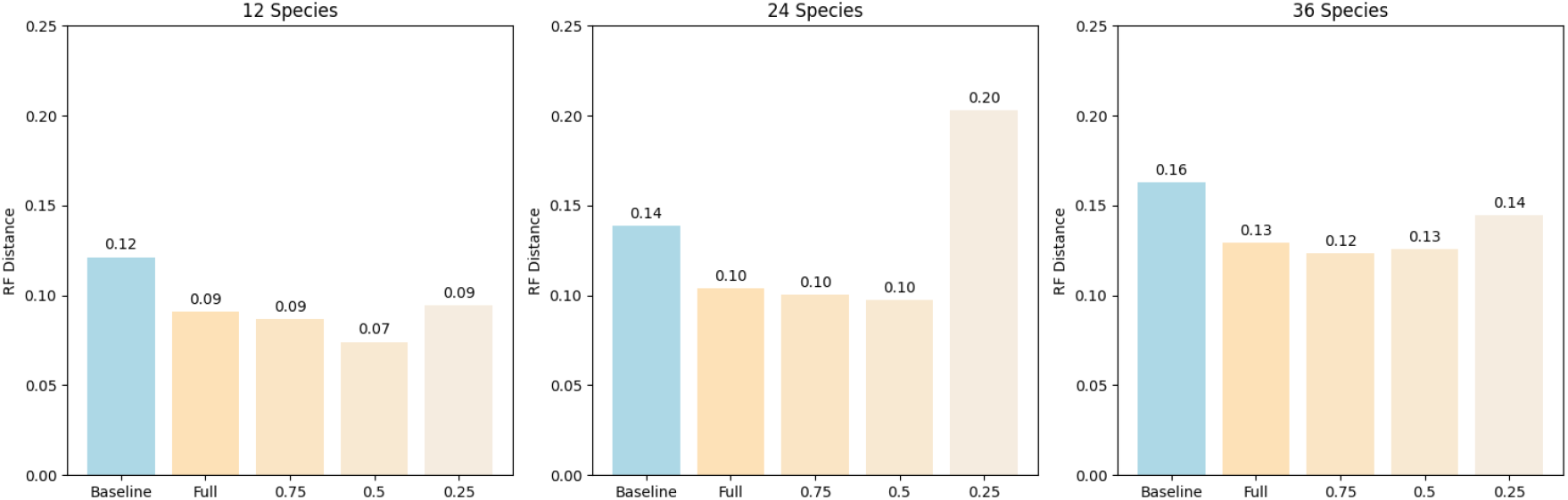
Reconstruction accuracy based on Quartformer predictions using wQFM.

### 4.2 Evaluation on Real–Biological Datasets

The approach of first using deep learning models to predict quartet topologies and then combining them into complete phylogenetic trees has been widely evaluated for real biological phylogenetic inference, as exemplified by Fusang [22]. However, a major limitation of deep learning models lies in their lack of interpretability, which remains a critical concern in the field of phylogenetics, where methodological rigor and reliability are of paramount importance. In contrast, traditional methods for inferring phylogenetic trees from molecular data, such as NJ, ML, and BI, are built upon solid statistical foundations. These classical methods search for phylogenetic trees by optimizing explicit objective functions, and they provide well-defined parameter estimates and measures of uncertainty.

For testing on real biological datasets, we downloaded 19,444 phylogenetic trees from the Pfam database and extracted subtrees with a specific number of species. These subtrees were then used as input for the Indelible software to generate corresponding MSAs. To account for insertions and deletions in evolutionary processes, we incorporated simulation of these events into the original setting. Specifically, we set the length of the ancestral sequences to 1000 bp, with insertion and deletion rates both set to 0.01. The lengths of insertions and deletions followed a Zipf distribution with *α* = 1.5, with a maximum length of 50 bp. Finally, quartets and subtrees with specific numbers of leaves were induced to form the training data for both models.

To assess the reliability and effectiveness of deep learning model predictions, we employed likelihood as an evaluation criterion for the predicted topologies. The test dataset is based on the real biological data released by Azouri et al. (2024) [2], which contains MSAs of 7, 12, 15, and 20 species. For each MSA, tree topologies were inferred using three methods: RAxML, the baseline framework, and the Quartformer framework. The likelihood of each inferred topology was then calculated using RAxML to ensure a fair comparison. As shown in Figure 18, in most cases, the tree topologies inferred by deep learning models performed comparably to those obtained via RAxML under the likelihood–based criterion.

**Figure 18:**
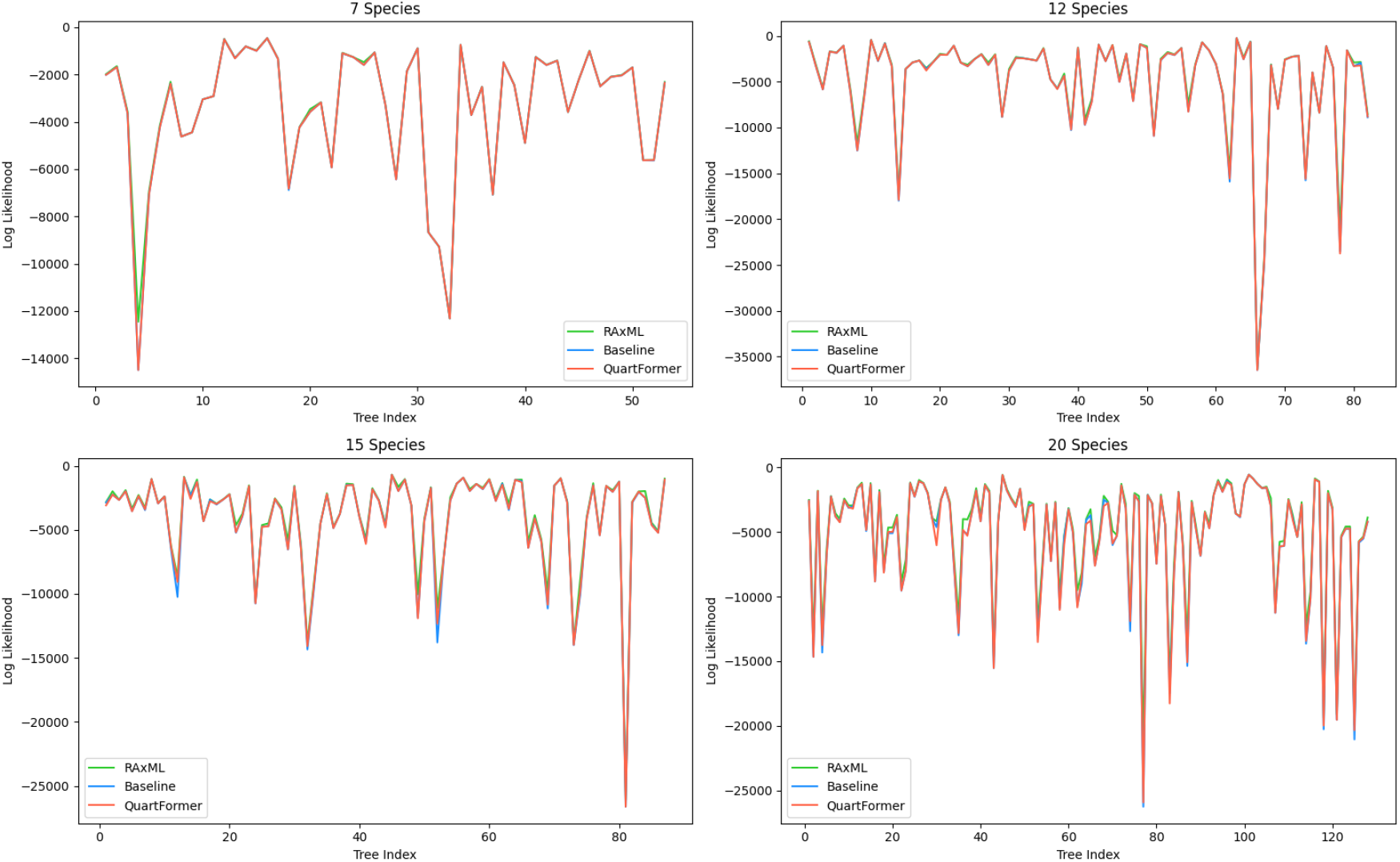
Likelihood comparison of trees inferred by RAxML, the baseline, and Quartformer.

Furthermore, we compared the likelihood of the baseline framework and the Quartformer framework across all test samples. As shown in Figure 19, For datasets with 7, 12, 15, and 20 species, the Quartformer framework outperformed the baseline framework, demonstrating a greater number of samples with higher likelihood values.

**Figure 19:**
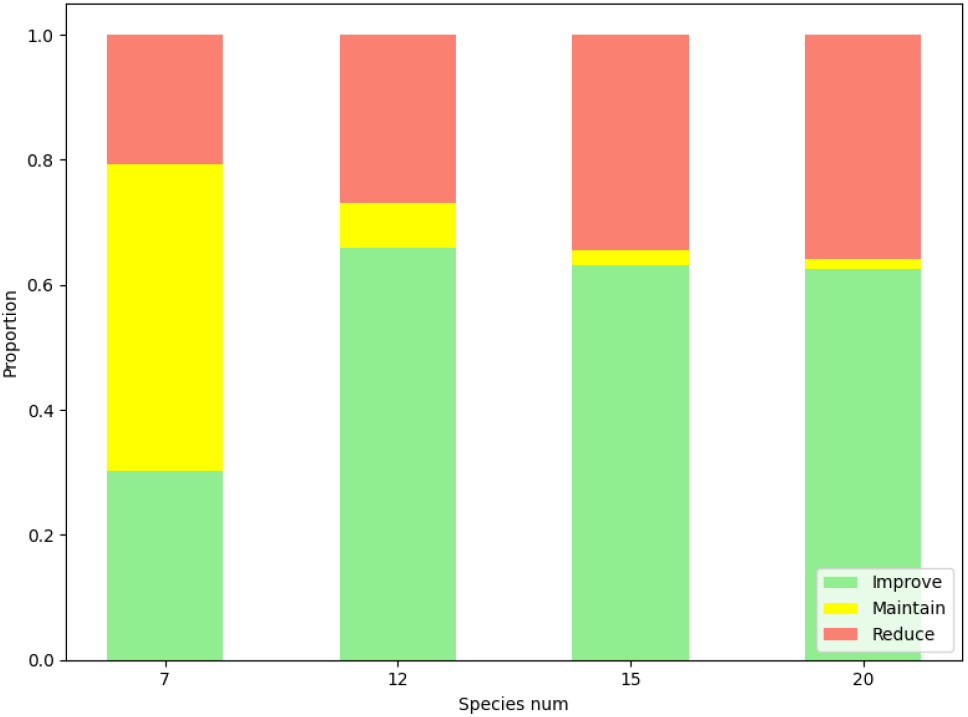
Likelihood comparison between baseline and Quartformer.

## 5. Theoretical Analysis of Quartformer’s Advantages

We reframe the training process of baseline and Quartformer within the framework of supervised learning. For simplicity, we assume that the training dataset is induced by a phylogenetic tree 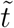. Let 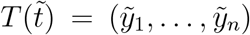, where 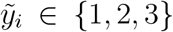 represents the quartet topology induced by 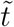, and *Q* = *{q*_1_, …, *q*_*n*_*}* denotes the set of MSAs corresponding to all quartets. Since our task is to predict 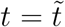 based on *Q*, the corresponding distribution over the training sample can be defined as:

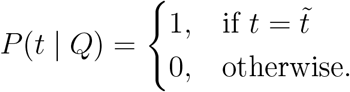

Given that 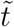 and 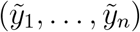 are in a one–to–one correspondence, this distribution can equivalently be rewritten as:

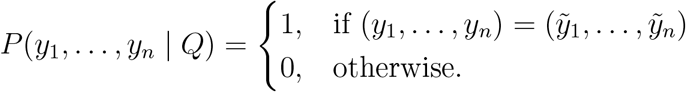

However, due to the (*y*_1_, …, *y*_*n*_) space having 3^*n*^ possible combinations, it is unrealistic to have a model directly predict the distribution over this space for classification. Therefore, both models instead focus on fitting the marginal distribution of each quartet individually, effectively decomposing the original sample 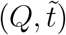 into *n* separate sub-samples. A key difference between the two frameworks lies in whether 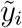 is determined by the individual *q*_*i*_ (as in the baseline) or by the entire *Q* (as in Quartformer). For each sample, our task is to infer the most likely quartet topology induced by the phylogenetic tree that best explains *Q*. From a theoretical standpoint, the optimal explanation naturally supports Quartformer’s assumption. In contrast, the baseline assumes that 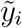 is entirely determined by *q*_*i*_, ignoring the global constraints imposed by other quartets, which often leads to inconsistent predictions. Therefore, we advocate representing each sample as 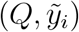.

We denote the baseline model as **f**_*B*_(*y*_*i*_ |*q*_*i*_; *θ*_*B*_), where *θ*_*B*_ represents the trainable parameters of the baseline model. Similarly, the Quartformer model is defined as

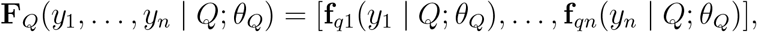

where *θ*_*Q*_ denotes the trainable parameters of the Quartformer model. The loss functions for both models are:

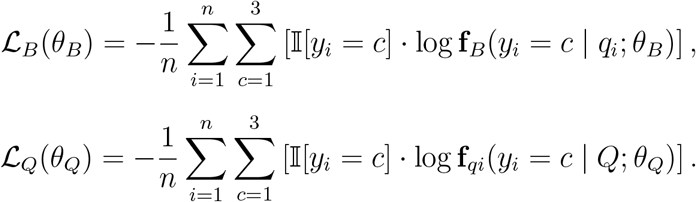

Taking the difference between these losses:

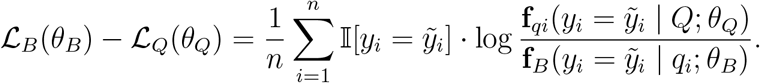

Since 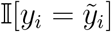 represents the marginal distribution *P*_*i*_(*y*_*i*_ | *Q*), we can rewrite this as:

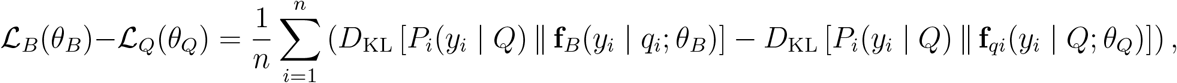

where *D*_KL_(·∥·) denotes the KL divergence between two distributions. During training, *P*_*i*_(*y*_*i*_ |*Q*) and **f**_*B*_(*y*_*i*_ |*q*_*i*_; *θ*_*B*_) differ structurally: the latter is a local model based solely on the individual MSA *q*_*i*_, while the former reflects the true conditional distribution that captures global dependencies among all quartets in *Q*. As a result, **f**_*B*_ (*y*_*i*_ |*q*_*i*_; *θ*_*B*_) lacks the global information necessary to model inter–quartet interactions, leading to a less faithful representation of the underlying distribution. From the perspective of model structure, **f**_*B*_(*y*_*i*_ |*q*_*i*_; *θ*_*B*_) can be viewed as a basic component within **f**_*qi*_(*y*_*i*_ |*Q*; *θ*_*Q*_). This implies that Quartformer generalizes the baseline model and, at minimum, retains its fitting capacity—if not surpassing it—by further incorporating global quartet-level dependencies. Thus, theoretically, ℒ_*B*_(*θ*_*B*_) *≥* ℒ_*Q*_(*θ*_*Q*_) when training converges, indicating that Quartformer has stronger fitting capabilities and can better approximate the target distribution. This explains Quartformer’s significant performance gains on simulated datasets. However, on real biological datasets, while Quartformer generally outperforms baseline, the improvements are less pronounced. We attribute this mainly to the fact that the tree topologies in the training data and their simulated MSAs do not fully conform to the ML criterion. This discrepancy is also evident in the tree construction tasks using RAxML in the simulation experiments, where even on small–scale phylogenetic reconstruction tasks, the tree topology corresponding to the ML still differs from the true topology used to simulate the MSA.

## 6. Conclusion and Discussion

In this study, we propose a deep learning–based framework for phylogenetic tree construction called Quartformer. Specifically, Quartformer builds upon the previous framework of independently predicting quartet topologies and then combining them, by incorporating a sparse attention model. This model effectively integrates information from different quartets within the same phylogenetic tree, thereby improving the consistency of predictions across all quartets. Through experiments on simulated datasets and theoretical analysis, we demonstrate that the introduction of the sparse attention model significantly enhances the learning capability of the deep learning framework. Furthermore, we provide a detailed analysis of the framework’s inference speed and accuracy. Finally, we test the framework on real datasets, and the results show that its predictions are largely consistent with the ML criterion, further validating its reliability in handling real biological data. The sparse attention model not only improves the accuracy of quartet topology inference but also provides new insights into integrating deep learning models into other subtree combination algorithms.

Although the deep learning model’s strong fitting ability has been fully validated on simulated datasets, and the framework performs even more prominently after the introduction of the sparse attention component, its reliability when applied to real biological scenarios still faces some challenges. The current deep learning model is primarily trained on simulated datasets, but real datasets are typically more complex and diverse, which poses a challenge to the model’s generalization ability. For instance, in testing on real datasets, using likelihood values as a criterion, the likelihood values of the deep learning model’s outputs still show some disparity when compared to those of RAxML, and the performance improvement of Quartformer is smaller than that observed on simulated datasets. This suggests that the training samples generated by the simulated data deviate from the ML standard. This issue may potentially be addressed by exploring the following two approaches. The first is the design of the training dataset, such as including reliable real phylogenetic trees or trees estimated using the ML method in the simulated dataset, while also considering the comprehensiveness of the samples and increasing the number of training samples that are difficult to classify with conventional methods. The second approach involves the design of the training algorithm, using regularization techniques to prevent the model from overfitting to the noise or specific patterns in the training data.

Another constraint for applying deep learning algorithms to real tasks is the inference speed. Expanding the model to handle larger species counts for topology estimation is an important direction for future research. The number of quartets grows at a rate of O(*n*^4^) with the increase in species count, and when the number of species becomes too large, predicting all quartets becomes impractical. This study demonstrates the feasibility of predicting only a subset of quartets and combining them, and in theory, this method can be further expanded by integrating subtree combination algorithms such as Matrix Representation with Parsimony (MRP) [14]. However, its accuracy still requires experimental validation. Additionally, improving the model’s ability to handle longer sequences is an urgent issue. The current method slices the original MSA data into fixed–length segments for input, which causes the model’s running time to grow nearly linearly with the DNA sequence length. However, in real biological applications, genomic–level input data is often encountered, which puts significant pressure on the deep learning model’s runtime and storage requirements. One feasible solution is to adopt the “Bag–of–Words Frequency Embedding” strategy proposed by Berk Alp Yakici [23], which compresses MSA quartets of any length into fixed–length vectors for input to the quartet classification model. This approach will significantly improve the deep learning model’s inference speed and practicality.

## 7. Code Availability

The source code related to this work is available at: https://github.com/thedata-learner/Quartformer. Pfam family trees used in this study can be downloaded from the following database: https://ftp.ebi.ac.uk/pub/databases/Pfam/releases/Pfam35.0/trees.tgz.

